# Glycosylated Antibiotics: New Promising Bacterial Efflux Pumps Inhibitors

**DOI:** 10.1101/2021.03.31.437923

**Authors:** Zahraa Ali Kamaz, Haruna Isiyaku Umar, Parth Doshi, Praveenya Suri

## Abstract

**Background:** Antimicrobial resistance is considered a major concern problem; bacteria have evolved mechanisms to overcome antibiotics’ action through evolutionary process. One main resistance mechanism that bacteria developed is the pumping of the antibiotics out of bacterial cells by transmembrane transporter proteins known as efflux pumps.

**Materials and methods:** To overcome bacterial resistance guided by efflux pumps, efflux pumps inhibitors (EPIs) are small molecules that obstruct efflux pumps binding sites and its structural assembly leading to disability in the efflux pumps normal function, new EPIs which under the current study are created by modifying the chemical structure of most common antibiotics including Ampicillin, Penicillin, Chloramphenicol, Ciprofloxacin and Tetracycline, such antibiotics are modified by adding N-acetyl glucose amine moiety to acceptor OH group of the respective antibiotic, the newly modified antibiotics are glycosylated EPIs. To test the effectiveness of the new EPIs in inhibiting AcrB-TolC and MexA-OprM efflux pumps functions, ADME properties for all of glycosylated antibiotics have been measured through applying Lipinski’s role of 5, docking and simulation studies have been included as well.

**Results:** Docked glycosylated tetracycline has given the highest binding energy in the active sites of both pumps, with −9.4 against AcrB and −8.8 against MexA. The simulation study has confirmed the binding of the glycosylated tetracycline in the active sites of both pumps, as well as its stability during the biological dynamicity of both pumps (opening and closing channels).

**Conclusion:** The results validation requires a long simulation time about 50 ns or more which was un applicable due to cost limitation, however, the newly glycosylated antibiotics have promising results that might make it eligible as drug candidates to overcome bacterial resistance.

## Introduction

Bacterial antibiotics resistance is a globally concern problem, bacteria have evolved mechanisms to counter antibiotics’ actions, main one involving the pumping out of antibiotics from bacterial inside into the external environment by specialized membrane proteins known as efflux pumps, which confer bacterial resistance to a wide range of antibiotics [1,2]. Efflux pumps are bacterial transport proteins that extrude toxins and antibiotics out of bacterial cells by energy dependent mechanisms [3], thereby classified into ABC active transporters, derived their energy from ATP molecules while other transporters’ families derived their energy by chemical gradients which are MFS (Major Facilitator Superfamily), MATE (Multidrug and toxic compounds extrusion), SMR (The small multidrug resistance), RND (Resistance Nodulation-division), and DMT (Drug metabolite transporter) [4,5]. Well studied efflux pumps are AcrB-TolC in *E.coli* and Mex-OPrM that mostly found in *Pseudomonas aeruginosa* [6,7], molecular based structures for both transporters have been elucidated extensively, TolC and OprM are tunnel structures that span the outer membrane of Gram-negative bacteria, It’s a homotrimer composed of α- helical barrel, β-barrel and α/β mixed fold [1], AcrB and Mex are fusion or linker proteins that are directly in contact with TolC and OprM respectively, whereas the inner cytoplasmic domain is the transporter protein, some of its parts spanning the cytoplasmic membrane to come in contact with fusion parts, for better descriptions, both pumps structures are illustrated in **(Fgure 1)**, [1,8].

**Figure 1.**
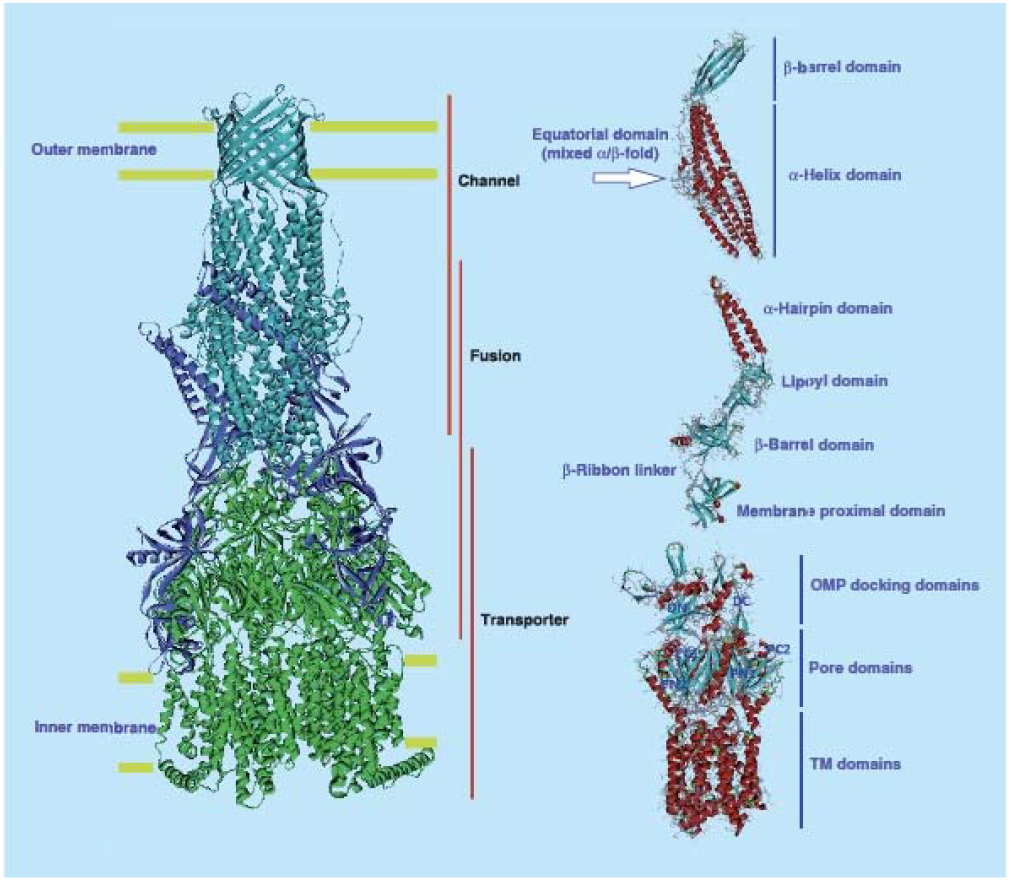
The structural assembly of MexA efflux pump. The structural components of MexA efflux pump on the left panel, that its similar to the structure of AcrB efflux pump on the right side, the Image has been taken from [1].

As the important role of bacterial efflux pumps in bacterial treatments’ failures and concurrent bacterial infections, numerous natural and synthetic compounds have been repurposed as efflux pumps inhibitors (EPIs), such compounds either cause conformational change in the structure of efflux pumps to inhibit its functional assembly or block the substrates binding site of the efflux pumps [9]. PAβN (Phenylalanine-arginine β-naphthylamide) is the first discovered inhibitor for Mex-OprM efflux pump that blocks the binding site of the pump resulting in an increase of the efflux sensitive antibiotic inside bacterial cells, Pyridopyrimidine derivative D13-9001 restore the antibiotic activity by competitive binding with the substrate binding site of AcrB-TolC and Mex-OprM efflux pumps [10]. Several other natural EPIs are plants alkaloids, polyphenols, quinolones derivatives and others, majority of them with unknown exact mechanisms of action [4,11].

EPIs are considered as potential antibiotics adjuncts to overcome multi-drugs resistance, working by combining EPIs with antibiotics course of treatments, or simply just modifying the antibiotics structure to create new EPIs or so called antibiotics analogues such as tetracycline analogue that recently known to restore tetracycline effectiveness in resistant bacteria [12]. Glycosylation of antibiotics is a natural structural modification process that mostly common among macrolides, bacterial glycosyltransferase transfers UDP-glucose moiety into antibiotic’s OH receptor and circumvents the antibiotics activity [13,14]. Glycosylated antibiotic can be synthesized through applying the natural glycosylation principle to the specified antibiotic, such new class of antibiotics might restore the antibiotic activity in the resistant bacteria by working as EPIs or antibiotics analogues. To test the hypothesis, sweet antibiotics have been docked against the active binding site of AcrB and MexA efflux pumps and molecular dynamic simulation has been applied to figure out the conformational changes of the mentioned efflux pumps upon sweet antibiotics binding.

## Results

### 1. Pharmacokinetic properties

Lead likeness properties for all five glycosylated antibiotics were measured, Lipinski’s role of 5 to predict drugs likeness properties for all new glycosylated antibiotics were followed, the drugs criteria are MW of the lead ≤ 500, less than 5 hydrogen donors, less than 10 hydrogen bond acceptor, Log P or lipophilicity not greater than 5 [15], the lead-likeness properties for the glycosylated antibiotics along with Lipinski’s number of violations are listed in (**Table 1)**. All the glycosylated antibiotics have shown 2-3 number of violations to Lipinski’s role, major one is the MWs for the most of newly antibiotics were larger than 500 except for ciprofloxacin and chloramphenicol.

**Table 1.**
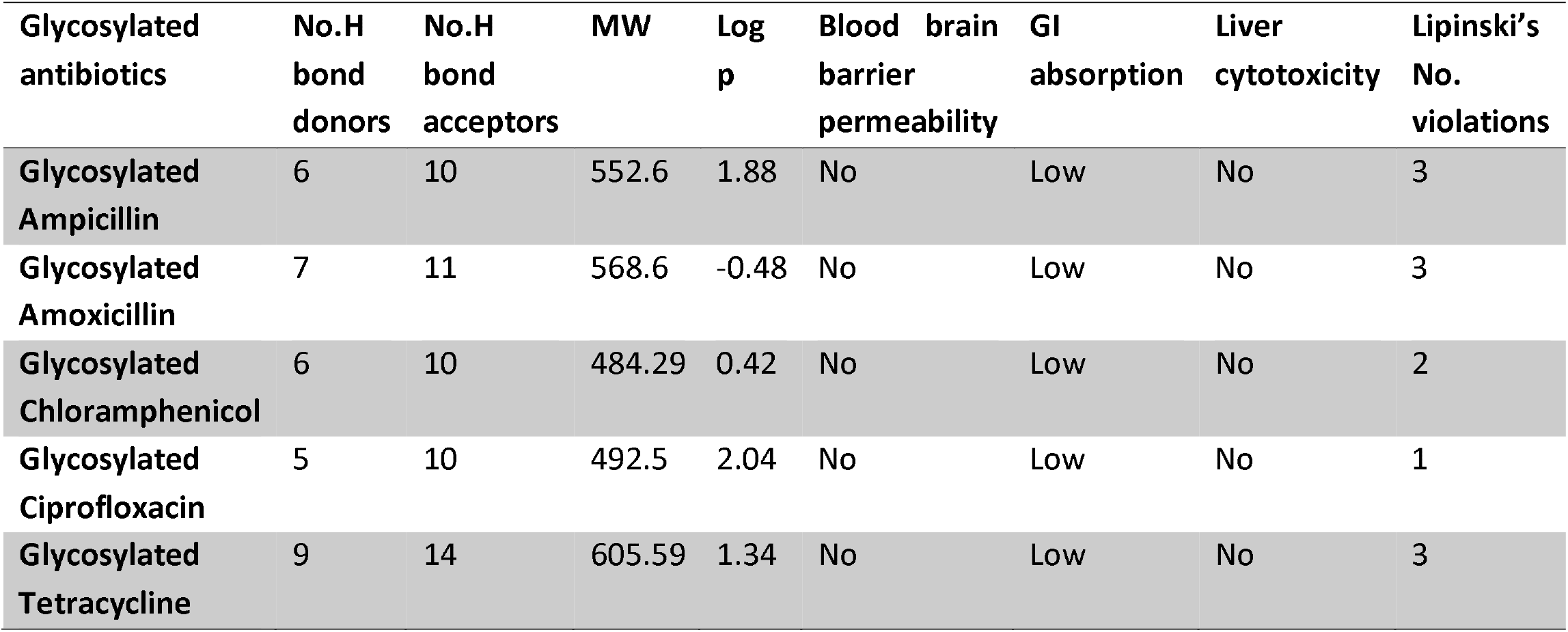
Pharmacokinetics properties of the newly glycosylated antibiotics.

### 2. Docking results

The binding energy for all ligands “glycosylated and non-glycosylated antibiotics” against AcrB and MexA efflux pumps are illustrated in (**Tables 2, 3)**. Glycosylated tetracycline has given the highest binding energy against both efflux pumps, with −9.7 against AcrB efflux pump and −8.8 against MexA, the interacted amino acids residues in the binding pockets for both pumps against glycosylated tetracycline are explained in **(Figures 2, 3)**. Glycosylated tetracycline has shown some similarity in the binding pocket of MexA efflux pump when compare it to the binding pocket of pyridopyrimidine derivative D13-9001 in the same pump, shared amino acids residues are GLN110, LYS 108 and SER 107 **(Figure 4)**. Two EPIs (PAβN and pyridopyrimidine derivative D13-9001) were included in the docking study against both efflux pumps for comparison.

**Table 2.** Docking results of N-acetyl glucose amine antibiotics against AcrB efflux pump in *E. coli* and MexA efflux pump in *Pseudomonas aeruginosa*.

| Ligand | Binding Energy (kcal/mol) |  |
| --- | --- | --- |
|  | ArcB | MexA |
| Glycosylated Amoxicillin | -8.7 | -7.5 |
| Glycosylated Ampicillin | -8.3 | -7.9 |
| Glycosylated Chloramphenicol | -8.0 | -7.8 |
| Glycosylated Ciprofloxacin | -9.0 | -7.8 |
| Glycosylated Tetracycline | -9.7 | -8.8 |

**Table 3.** Binding Energy of Non-glycosylated antibiotics docked against ArcB and MexA.

| Ligand | Binding Energy (kcal/mol) |  |
| --- | --- | --- |
|  | ArcB | MexA |
| **Amoxicillin** | -7.8 | -6.3 |
| **Ampicillin** | -7.7 | -6.7 |
| **Chloramphenicol** | -6.1 | -5.8 |
| **Ciprofloxacin** | -7.5 | -6.4 |
| **Tetracycline** | -8.3 | -6.8 |
| **PAβN** | -9.3 | -9.1 |
| **Pyridopyrimidine** | -8.1 | -9.6 |

**Figure 2.**
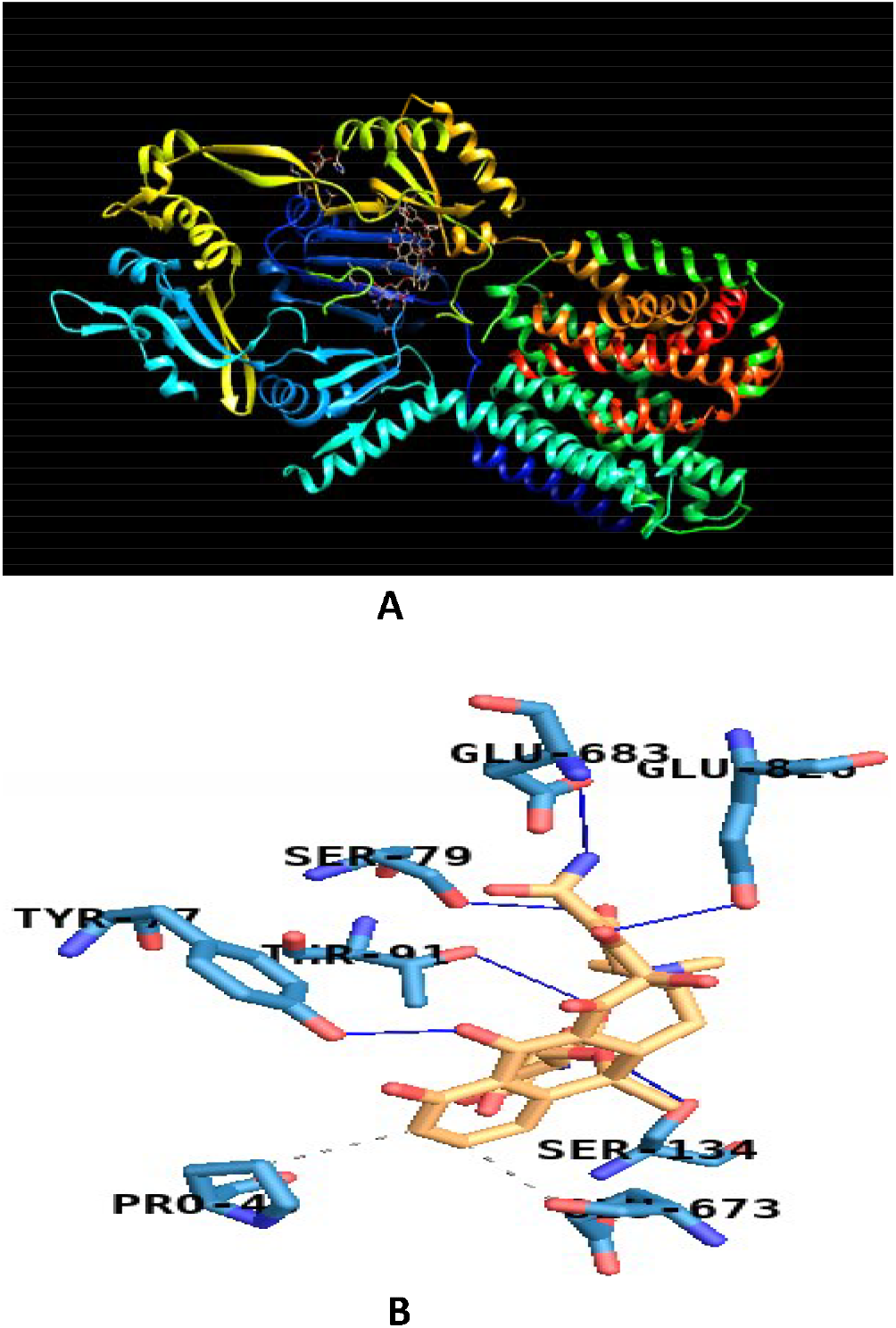
The binding site of the glycosylated tetracycline in Acrb efflux pump binding pocket. **A.** The left side is the binding pose of the glycosylated tetracycline **B.** The right side represents the interaction between amino acids residues of AcrB chain A and the glycosylated tetracycline, the new antibiotic showed hydrophobic interaction with PRO 40A, GLU 673A and Hydrogen bonds with TYR 77A, SER 79A, THR 91A, SER 134A, GLU 683A, GLU 826A.

**Figure 3.**
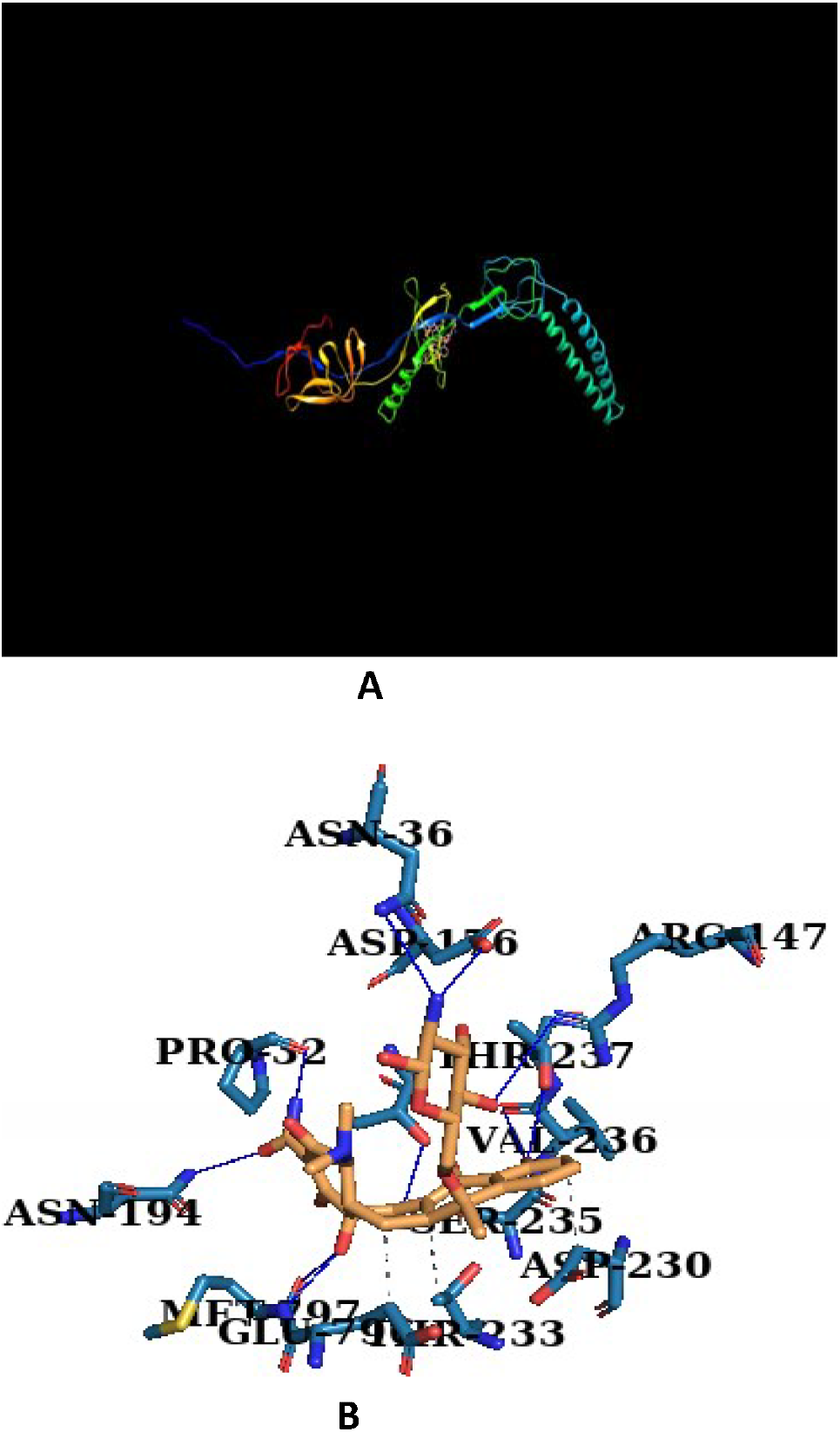
The binding sites of all of the glycosylated antibiotics in the binding pocket of MexA efflux pump. **A.** The left side, shows the binding pose of amoxicillin (cyan), ampicillin (pink), chloramphenicol (yellow), ciprofloxacin (red) and tetracycline (blue) in MexA. The drugs occupy similar region in MexA except amoxicillin. **B.** The right side is the interaction between residue of MexAB – OprM efflux pump and Glycosylated tetracycline, that showed hydrophobic interaction ASP 230D, THR 233D, GLU 796J and hydrogen bonds with PRO 32D, ASN 36D, ARG 147E, ASP 176D, THR 178D, ASN 194J, SER 235D, VAL 236D, THR 237D, MET 797.

**Figure 4.**
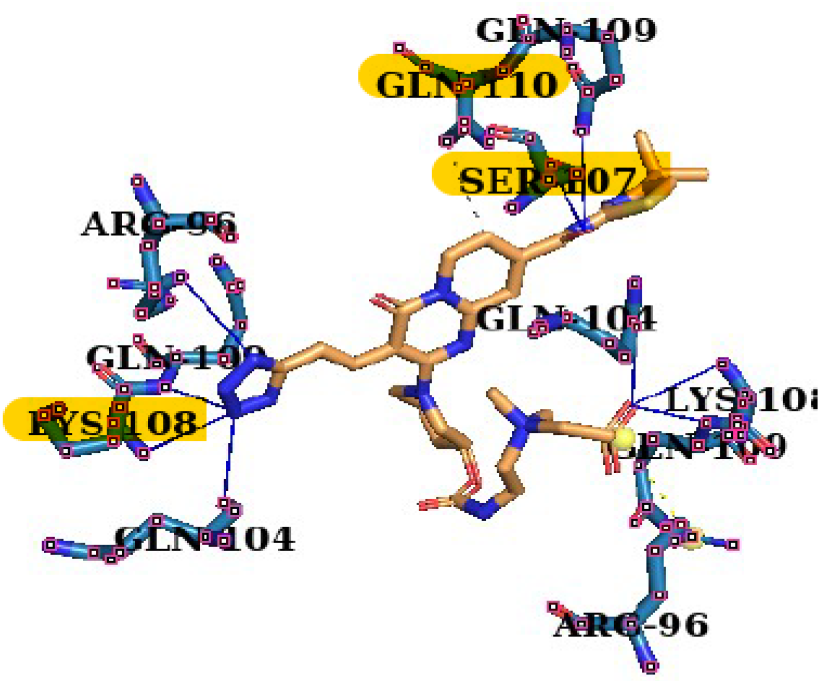
The binding site of pyridopyrimidine derivative D13-9001. Interacted amino acids residues in the binding pocket of MexA upon Pyridopyrimidine binding, shared residues with the glycosylated tetracycline binding pocket are highlighted in yellow.

### 3. Molecular dynamic simulation

RMSD value for the MexA-glycosylated tetracycline complex was plotted as in **(Figure 5)**, protein-ligand complex was not reaching the plateau since short time of simulation was devoted. However, the efflux pump dynamicity has been captured during the opening and the closing state of the pump **(Figure 6)**. The md simulation of AcrB efflux pump was done in 2 ns scale, RMSD value for AcrB-glycosylated tetracycline complex is explained in **(Figure 7)**, whereas **(Figure 8)** shows the complex conformational change during the opening and the closing process. The glycosylated tetracycline has given strong binding stability in the substrate binding sites for both studied pumps even during the biological dynamicity of the transporters pumps, the simulation results confirm the ability of the new antibiotic analogue to inhibit the binding site of AcrB and MexA efflux pumps.

**Figure 5.**
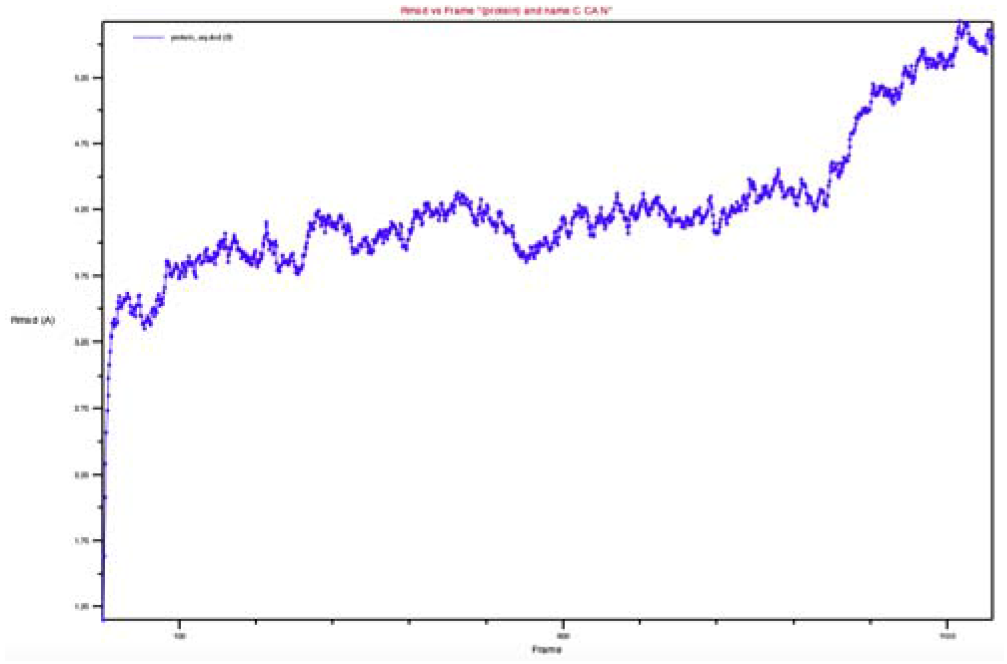
RMSD analysis for MexA-OprM-glycosylated tetracycline complex. CDC trajectory file for the solvate complex was used to analyze the RMSD value, 1150 frames were used, the protein-ligand complex didn’t reach the plateau because of short simulation time.

**Figure 6.**
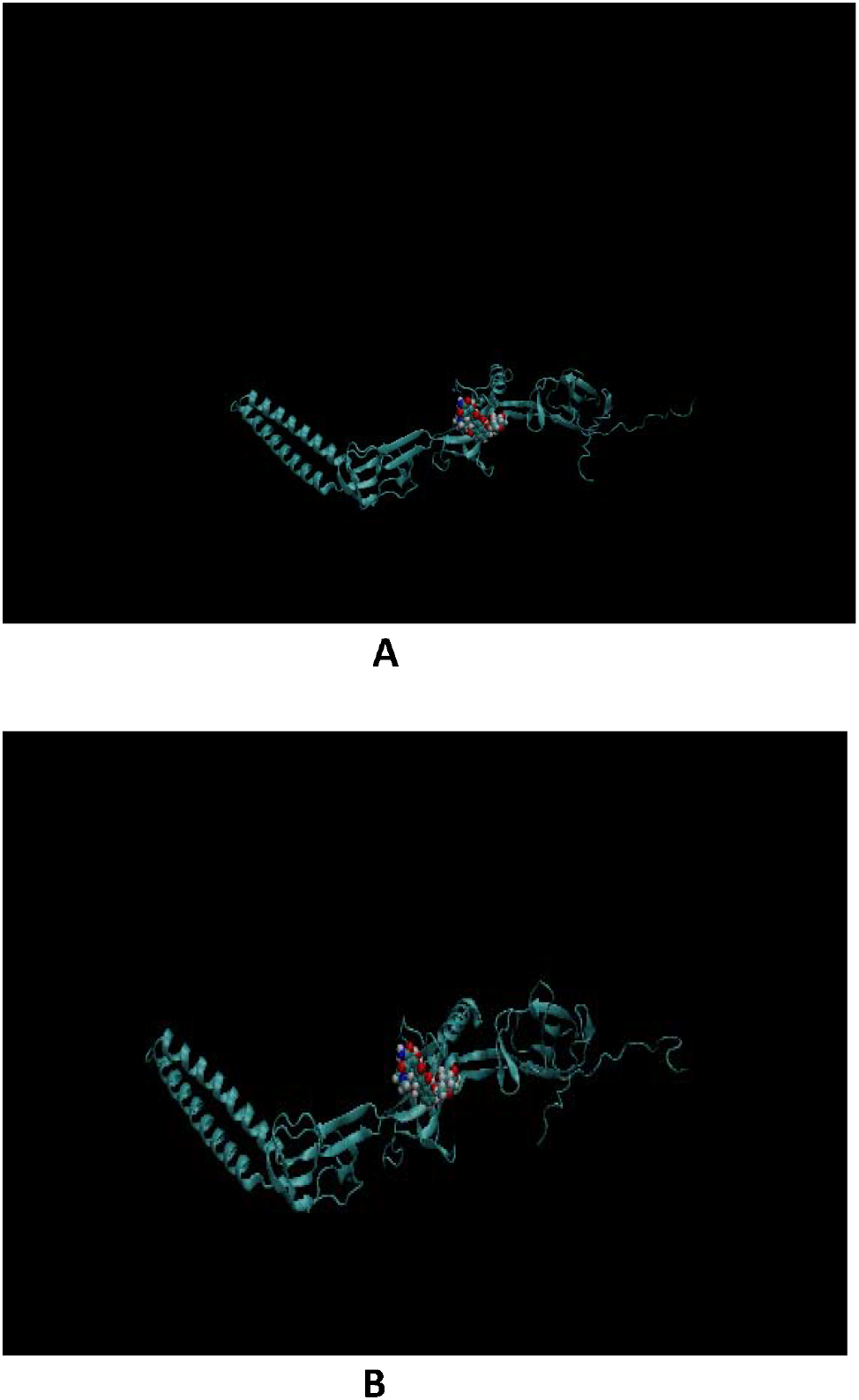
MexA efflux pump opening and closing states upon glycosylated tetracycline binding. the dynamic movement of MexA efflux pump upon binding with the glycosylated tetracycline, it shows the stability of the ligand during the closing of the channel **A.** and the opening of the channel **B.**

**Figure 7.**
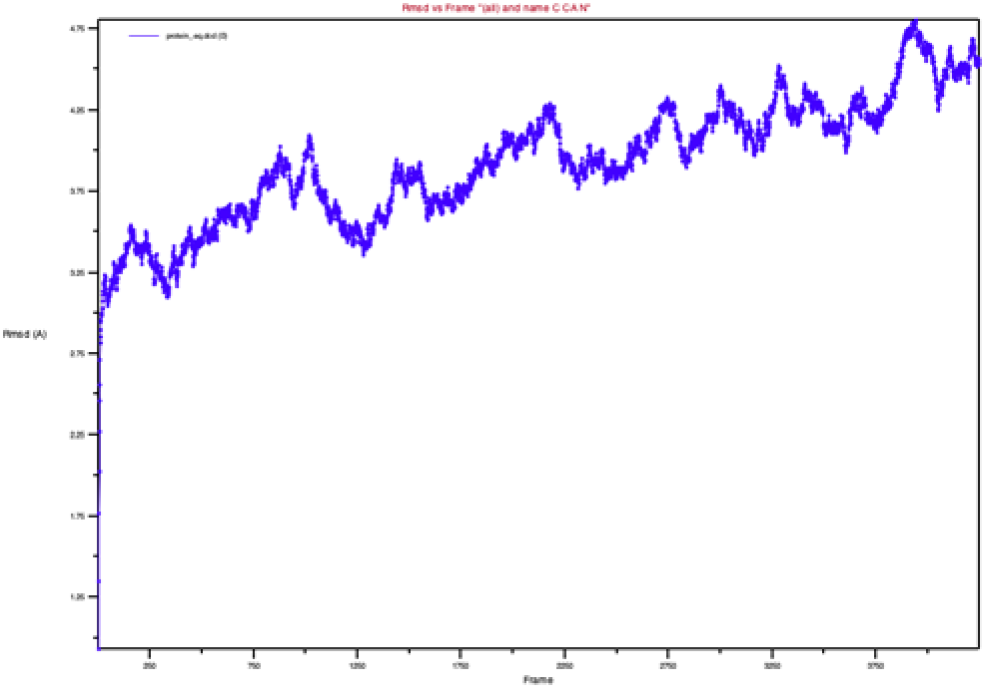
RMSD analysis for AcrB-glycosylated tetracycline complex. RMSD value for AcrB-glycosylated tetracycline was calculated using VMD program, 4250 frames were being used in the analysis of the CDC trajectory file. Again protein-ligand complex didn’t reach the plateau.

**Figure 8.**
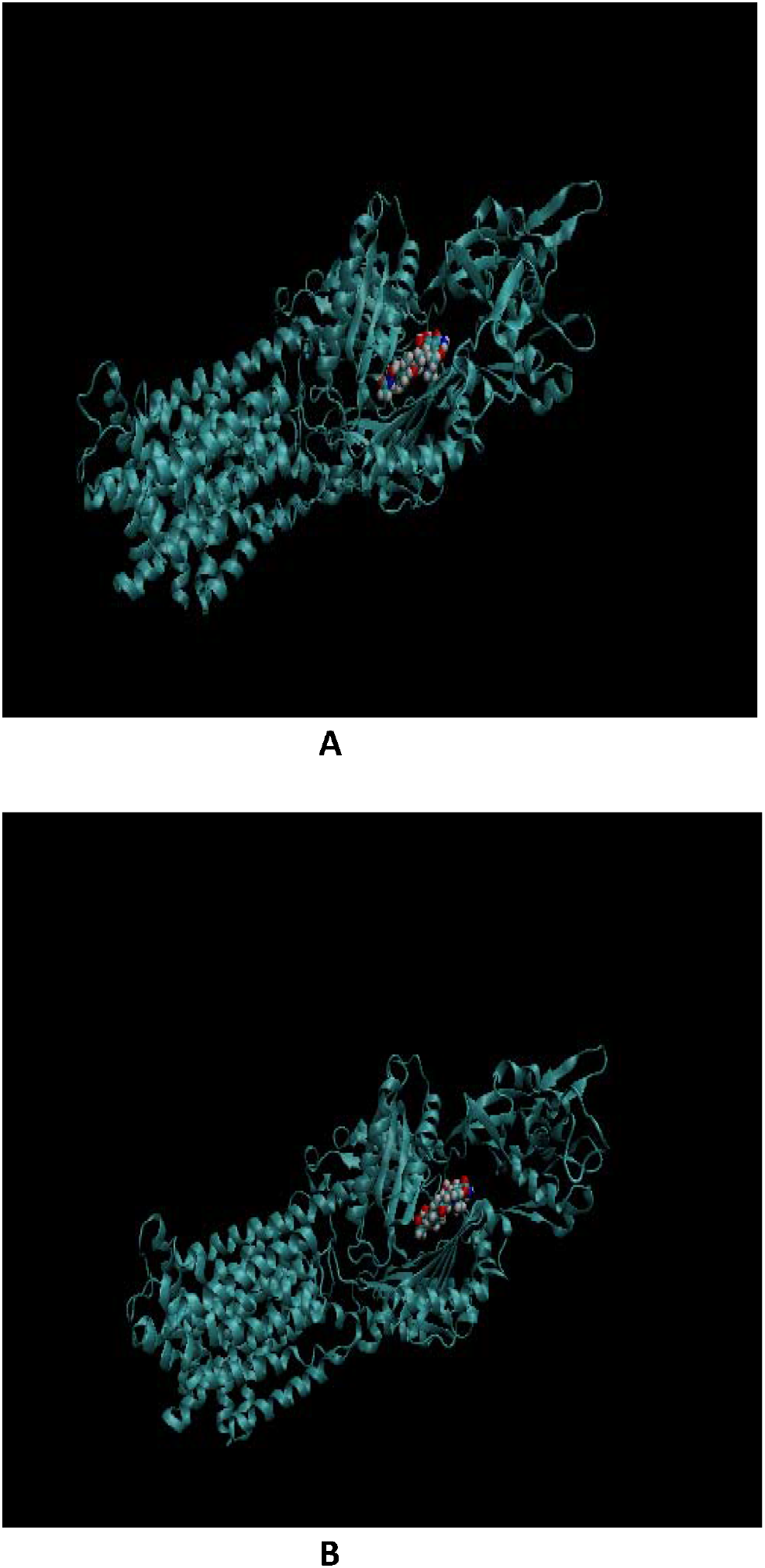
AcrB efflux pump opening and closing state upon glycosylated tetracycline binding. Shows the closing state **A.** and the opening of AcrB efflux pump **B.**, the glycosylated tetracycline is stable in the binding pocket of the target protein.

## Discussion

ADME properties are set of features that drugs candidates should meet to pass clinical trials, ADME stands for Absorption, Distribution, Metabolism and excretion (16), for the lead to be absorbed through GI tract and reached its target, it should has a good solubility and permeability (17). Since most of drugs are administered orally, so the oral bioavailability is such an important feature to test drug’s ability to reach its destination [18], Higher rate of drugs bioavailability is greatly dependent on its molecular weight (19). Three out of the five glycosylated antibiotics in the current study were seen to violate Lipinski’s rule for molecular weight determinant except for ciprofloxacin and chloramphenicol. As the importance of low molecular weight for orally administered drugs [20], however, there are other important features that all the understudied glycosylated antibiotics have such as, no toxicity and no blood brain barrier permeability. Furthermore, a study has confirmed the presence of many FDA approved antibiotics and antifungals that have violated the rule of 5 for low MWs but still are orally bioavailable because they have a specific group that works as a transporters’ substrate [21]. The rule of 5 appears as important data set that the orally- rout drugs should meet, in a study has examined about 111 non-oral administered drugs, concluded that Lipinski’s criteria for such drugs are not so important to meet [22].

The drugs’ journey starts by its absorption through gastrointestinal system, reaching blood stream and binding with the specified target to exert its action. The targeted proteins are two bacterial efflux pumps known as AcrB and MexA, the glycosylated tetracycline has shown the highest binding energy against both target proteins. The highest binding energy with the targeted proteins which are AcrB and MexA in the current study, the more drugs stability and effectiveness (23). Tetracycline resistance mediated by efflux pumps has increased recently, various factors are responsible for such resistance rise including the pumps genes are DNA mobilized elements that can transfer from one bacterium to another (24). Another factor is the long-term use of tetracycline causing the bacteria to co-evolve itself to extrude such toxins outside of the bacterial cells [25]. Developing molecules that target bacterial efflux pumps has been considered as a new treatment approach, one of the big issues that hurdle the use of EPIs is their toxicity to human cells [26]. Finding safe and effective EPIs is on the increase, of which one potential way is through modifying the structure of the effluxed antibiotics, such modification might help in blocking the binding capability of the effluxed antibiotic and successfully reverse the antibiotic resistance mechanism [27]. New tetracycline derivative known as glycylcycline has been developed as a combination therapy along with tetracycline to overcome Tet efflux pump resistance [27]. Glycylcycline which sometimes known as a tigecycline has similar structure to tetracycline with substitution of an N-alkylglycylamido group at D-9 position [28]. Another modification can be achieved through adding various glycosidic chains at C, O, N atoms on antibiotics, such glycosylation is a naturally occurring process that have been observed among number of antibiotics including macrolides and streptomycin [29]. Exploiting the glycosylation principle in creating new EPIs has shown promising results in compare to synthetically available EPIs. Because the targets are efflux pumps transporter proteins which are dynamic structures, molecular dynamic simulation has been applied to test the stability of the newly drugs in the binding pockets of its targets.

The molecular dynamic simulation results showed the glycosylated tetracycline stability in the binding sites of the studied efflux pumps, the efflux pump dynamicity has been captured during the opening and the closing state and the RMSD values for protein-ligand complexes were plotted. The molecular dynamic simulation studies of bacterial efflux pumps are a new era of interesting, investigating ligands stability among different conformational changes that efflux pumps undergone such as the transition from substrate binding state into opening state is facing issues at large scale proteins level [30]. The first md simulation studies for AcrB-TolC and MexA-OprM have been started since 1999 and still ongoing progresses till now, different simulation approaches have been used including mutable basin simulation, steered md simulation and unbiased equilibrium md simulation [31]. Identifying ligand binding site or what’s so called the hotspots is crucial in drug discovery, mixed solvent md simulation is successfully used for such a purpose, one of its big limitation is the cost of running such kind of simulation [32].

### Conclusion

Glycosylated tetracycline is showing promising results to work as EPIs without worrying about safety issues, however, it’s clinical approving require applying longer simulation time of 50 ns or more, as well as further in vitro studies are required to measure its activity against resistant bacteria. Glycosylation of other antibiotics with lower molecular weight and less Lipinski’s role of 5 violations are recommended.

## Materials and Methods

### 1. Glycosylation of antibiotics

Three dimensional structures of amoxicillin, ampicillin, chloramphenicol, ciprofloxacin and tetracycline were retrieved from pubchem web server (https://pubchem.ncbi.nlm.nih.gov/) in simple document format (SDF). The Pubchem CIDs for the studied antibiotics are 33613, 6249, 5959, 2764, and 54675776 respectively.

N-acetyl glucose amine moiety has been added to the hydroxyl acceptor group for each selected antibiotic, using Marvin sketch software 20.20, then the ligands were energetically optimized to the most stable structures using merk molecular force field 94 (MMFF94).

### 2. Pharmacophore kinetics properties

The drug-likeness properties have been measured for all glycosylated antibiotics using the website http://www.swissadme.ch/. The role of 5 for Lipinski as well as others features have been measured including the oral bioavailability, GI absorption, Blood brain barrier permeability, and liver cytotoxicity.

### 3. Molecular docking

3D x-ray crystallographic structures of AcrB and MexA were retrieved from the protein data bank (PDB) with IDs 2I6W and 6TA6 respectively. After which the proteins were prepared for docking and protein-ligand complexes were minimized using the relevant tools in Cresset Flare© software, version 4.0 (https://www.cresset-group.com/flare/). The protein minimization was done based on the GAFF force field, with gradient cutoff of 0.200 kcal/mol/A and iterations was set to 2000 iterations [33]. Furthermore, the docking was achieved using flexible docking protocol [34]. In brief; Python Prescription 0.8, a suite housing Auto Dock Vina, was employed for the molecular docking study of the selected ligands with the two efflux pumps. The specific target site for AcrB was set using the grid box with dimensions (16.94 × 4.797 × 11.6467) Å, while that of MexA was (95.452 × 111.512 × 34.633) Å and their centres were adjusted based on the active site of the enzymes. At the end of the molecular docking, 10 binding poses with least binding affinity for each protein-ligand complex were generated for all the ligands, scoring results were also created after retrieving from text files which were employed for comparative analysis. The protein-ligand complexes as well the molecular interactions were all visualized in 3D and 2D using PyMOL© Molecular Graphics (version 2.4, 2016, Shrodinger LLC) and Protein- Ligand interaction Profiler PLIP server (https://plip-tool.biotec.tu-dresden.de/plip-web/plip/index).

### 4. Molecular dynamic simulation

For glycosylated tetracycline which give the lowest binding affinity against both efflux pumps’ binding sites, molecular dynamic simulation was performed using NAMD 2.8, the topology file for the efflux pumps proteins were generated using automatic PSF builder in VMD program, whereas, the topology file for the ligand was generated using CHARMM GUI server (http://www.charmm-gui.org/), 2ns simulation was performed with time step of 2 fs, periodic boundary conditions were used, the temperature kept at 310 K and the pressure was stable at 1.013 bar, 100 steps of equilibration were completed before starting the solvate simulation, the whole md simulation protocol can be found [35]. CDC trajectory file for both efflux pumps-ligand complexes was analyzed in the form of measuring RMSD values using VMD graphical visualization program 1.9 version.

## Authors’ contributions

ZK carried out the study experimental design, molecular dynamic simulation work and writing the first draft of the research paper. HU carried out the modification of antibiotics’ structures work, did part of the docking work and reviewed the paper. PD did most of the docking work, and analyzed the docking results. PS confirmed the docking results by replicated it and reviewed the paper.

## Supplementary materials

**Figure 9.**
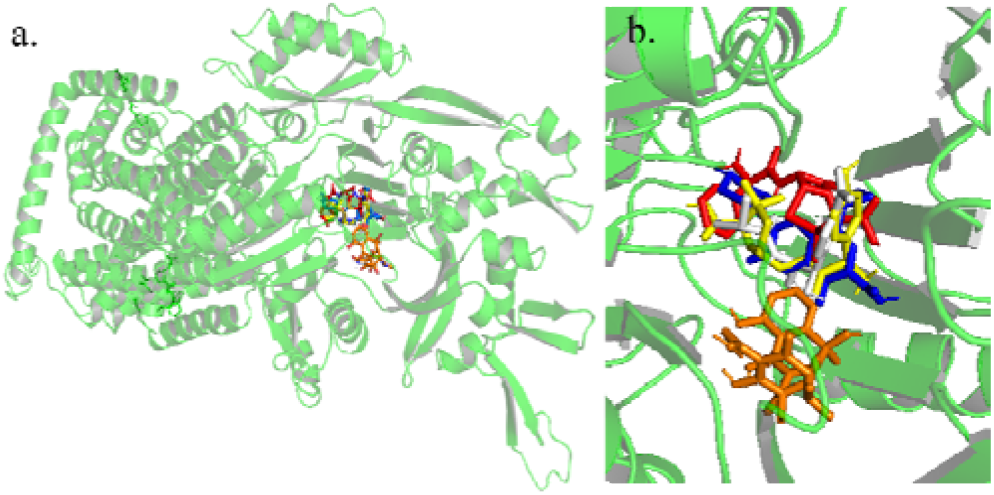
All non-glycosylated antibiotics binding with AcrB efflux pump. Binding pose of amoxicillin (yellow), ampicillin (red), chloramphenicol (white), ciprofloxacin (blue) and tetracycline (orange) in ArcB. The drugs occupy similar region in ArcB.

**Figure 10.**
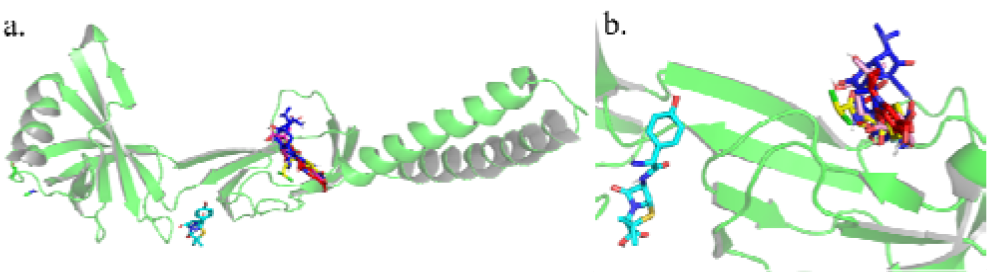
All non-glycosylated antibiotics binding with MexA efflux pump. Binding pose of amoxicillin (cyan), ampicillin (pink), chloramphenicol (yellow), ciprofloxacin (red) and tetracycline (blue) in MexA. The drugs occupy similar region in MexA except amoxicillin.

**Figure 11.**
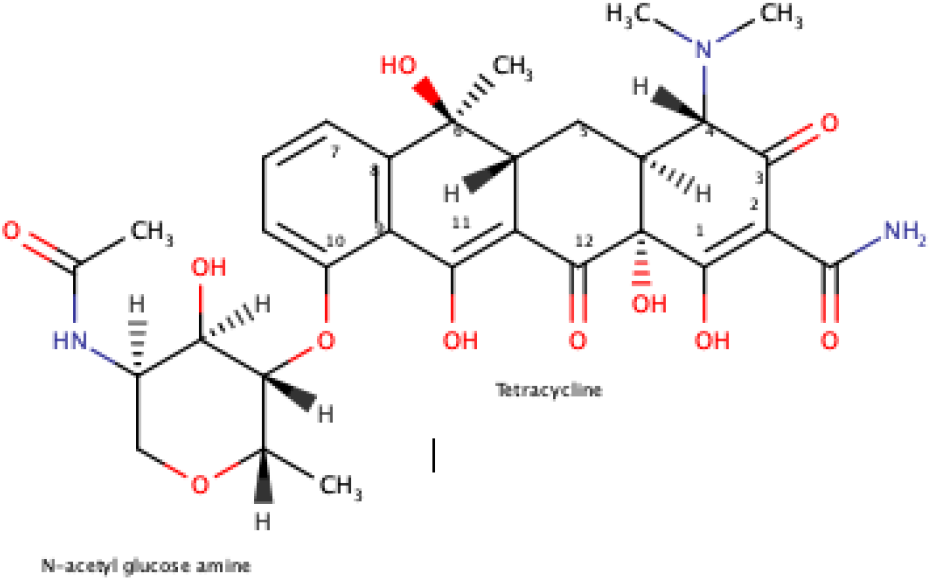
The chemical structure of the glycosylated tetracycline. N-acetyl glucose amine moiety is added to the hydroxyl group on atom 10 of tetracycline, all non-polar hydrogen atoms were removed and the structure is minimized to more stable state.

**Table 4.**
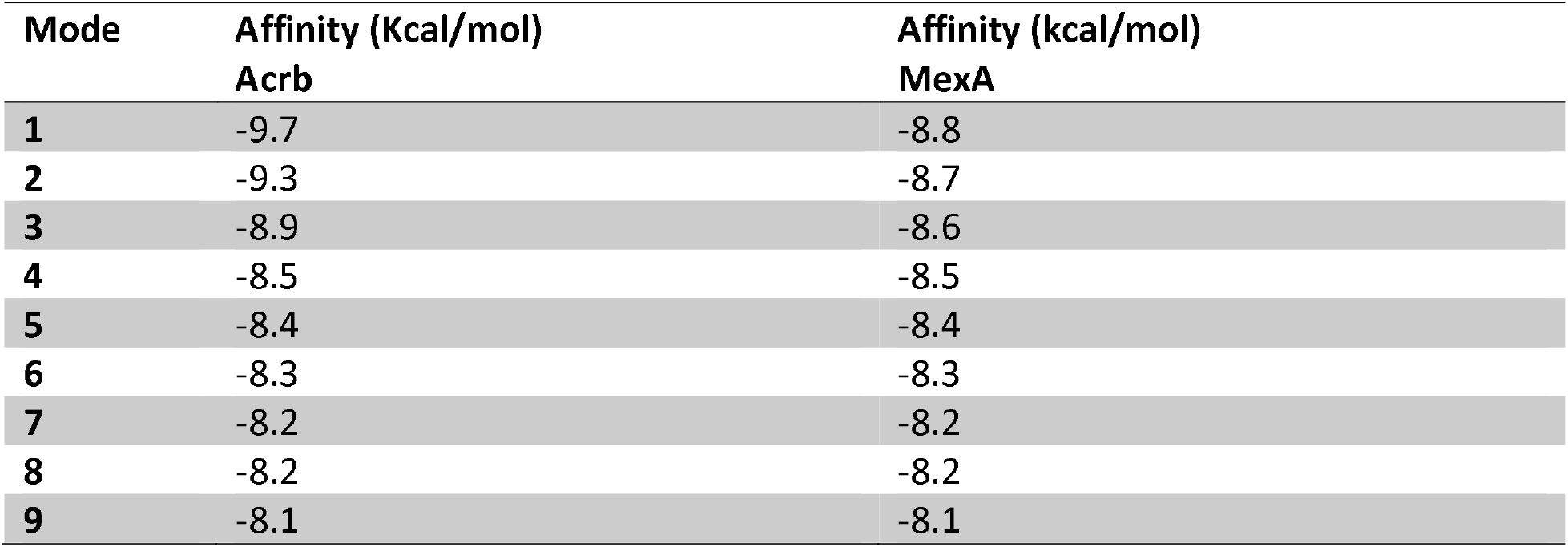
The binding affinities of the glycosylated tetracycline against AcrB and MexA efflux pumps.

| **Mode** | **Affinity (Kcal/mol)**<br>**Acrb** | **Affinity (kcal/mol)**<br>**MexA** |
| --- | --- | --- |
| **1** | -9.7 | -8.8 |
| **2** | -9.3 | -8.7 |
| **3** | -8.9 | -8.6 |
| **4** | -8.5 | -8.5 |
| **5** | -8.4 | -8.4 |
| **6** | -8.3 | -8.3 |
| **7** | -8.2 | -8.2 |
| **8** | -8.2 | -8.2 |
| **9** | -8.1 | -8.1 |

